# Predicting Brain Regions Related to Alzheimer’s Disease Based on Global Feature

**DOI:** 10.1101/2020.11.27.401950

**Authors:** Qi Wang, Siwei Chen, He Wang, Luzeng Chen, Yongan Sun, Guiying Yan

## Abstract

Alzheimer’s disease (AD) is a common neurodegenerative disease in the elderly, early diagnosis and timely treatment are very important to delay the course of the disease. In the past, most of the brain regions related to AD were identified based on the imaging method, which can only identify some atrophic brain regions. In this work, we used mathematical models to find out the potential brain regions related to AD. First, diffusion tensor imaging (DTI) was used to construct the brain structural network. Next, we set a new local feature index 2hop-connectivity to measure the correlation among different areas. The 2hop-connectivity utilizes the higher-order information of the graph structure relative to the traditional graph theory metrics. And for this, we proposed a novel algorithm named 2hopRWR to measure 2hop-connectivity. At last, we proposed a new index GFS (Global Feature Score) based on a global feature by combing 5 local features: degree centrality, betweenness centrality, closeness centrality, the number of maximal cliques, and 2hop-connectivity, to judge which brain regions are likely related to Alzheimer’s Disease. As a result, all the top ten brain regions in the GFS scoring difference between the AD group and the non-AD group were related to AD by literature verification. Literature validation results comparing GFS with local features showed that GFS outperforms individual local features. Finally, the results of the canonical correlation analysis showed that the GFS was significantly correlated with the scores of the mini-mental state examination (MMSE) scale and the Montreal cognitive assessment (MoCA) scale. So, we believe the GFS can also be used as a new index to assist in diagnosis and objective monitoring of disease progression. Besides, the method proposed in this paper can be used as a differential network analysis method in other areas of network analysis.

## 1 Introduction

Alzheimer’s disease (AD) is a common neurodegenerative disease in the elderly. It is a continuous process from the pre-clinical stage, mild cognitive impairment (MCI) to dementia. Effective intervention in the pre-dementia stage or MCI stage may delay or reverse the process of the disease. Therefore, early identification of AD patients in the pre-dementia stage or MCI stage, as well as early and timely intervention, are very important for the prognosis of patients. With the development of imaging technology, the detection of AD is not limited to the phenomenon of abnormal protein deposition. It may be an effective method for early diagnosis and monitoring of disease progression to analyze brain structural network information, such as brain connectome analysis (Fan et al., 2016).

A previous study (Liu et al., 2017) have shown that the change of topological feature of the brain structural network is a marker of many kinds of neuropsychiatric diseases. At present, there is some research work on brain structural networks based on graph theory (Sanz-Arigita et al., 2010; John et al., 2016). The common method is to analyze some local properties such as the degree centrality of nodes, clustering coefficient, and shortest path length of the brain structural network, and so on. Local features are difficult to reveal the whole characteristics of the network. Global property by combining local properties can reveal the topological characteristics of the network more effectively, but it is never easy for choosing which local indexes. In this paper, we first defined a new local feature index 2hop- connectivity of the network to analyze the brain network more completely.

In this work, 20 AD patients and 13 pre-dementia stages (non-AD) were recorded. We collected demographic data and clinical data, completed neuropsychological scale evaluation, and DTI scans. After image preprocessing, the brain structural network was constructed based on the number of fibers between different brain regions. The data of the AD group and the non-AD group were analyzed to get the local topological features of the brain structural network. At the same time, we designed an algorithm named 2hopRWR to get the local feature index 2hop-connectivity, and then we proposed a new index GFS by combing four classical local features and 2hop-connectivity. As a result, we predicted and analyzed the top 10 brain regions according to the GFS scoring difference between the AD group and the non-AD group. Then, we analyzed the correlation between the GFS and the cognitive scale scores by canonical correlation analysis (CCA). Finally, we discussed the strengths and limitations of our work and its prospects.

## 2 Materials and methods

### 2.1 Data collection and pre-processing

In this research, 20 AD patients and 13 healthy control (non-AD) were recorded. We collected demographic data and clinical data, completed neuropsychological scale evaluation (Supplementary materials table 1), and DTI scans. After image preprocessing, the deterministic fiber tracking FACT method was used to construct the brain structural network. We use the AAL brain atlas to divide each subject’s brain into 45 left and right symmetrical brain regions, 90 brain regions in total. Each node represents a brain region in the brain structural network. The fiber connection between any two brain regions is represented by an edge, and the edge weight represents the fiber number. The fiber number (FN) matrix of 90 brain regions was obtained by PANDA (Pipeline for Analyzing BraiN Diffusion imAges) (Cui et al., 2013).

**TABLE 1.**
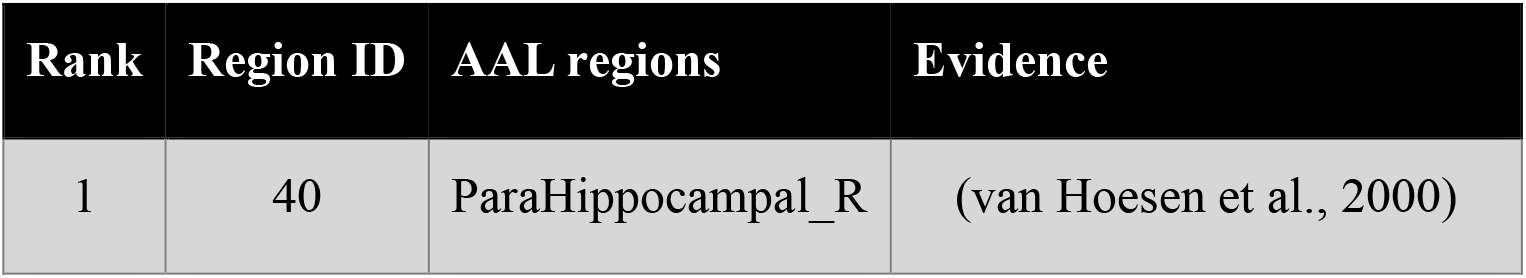
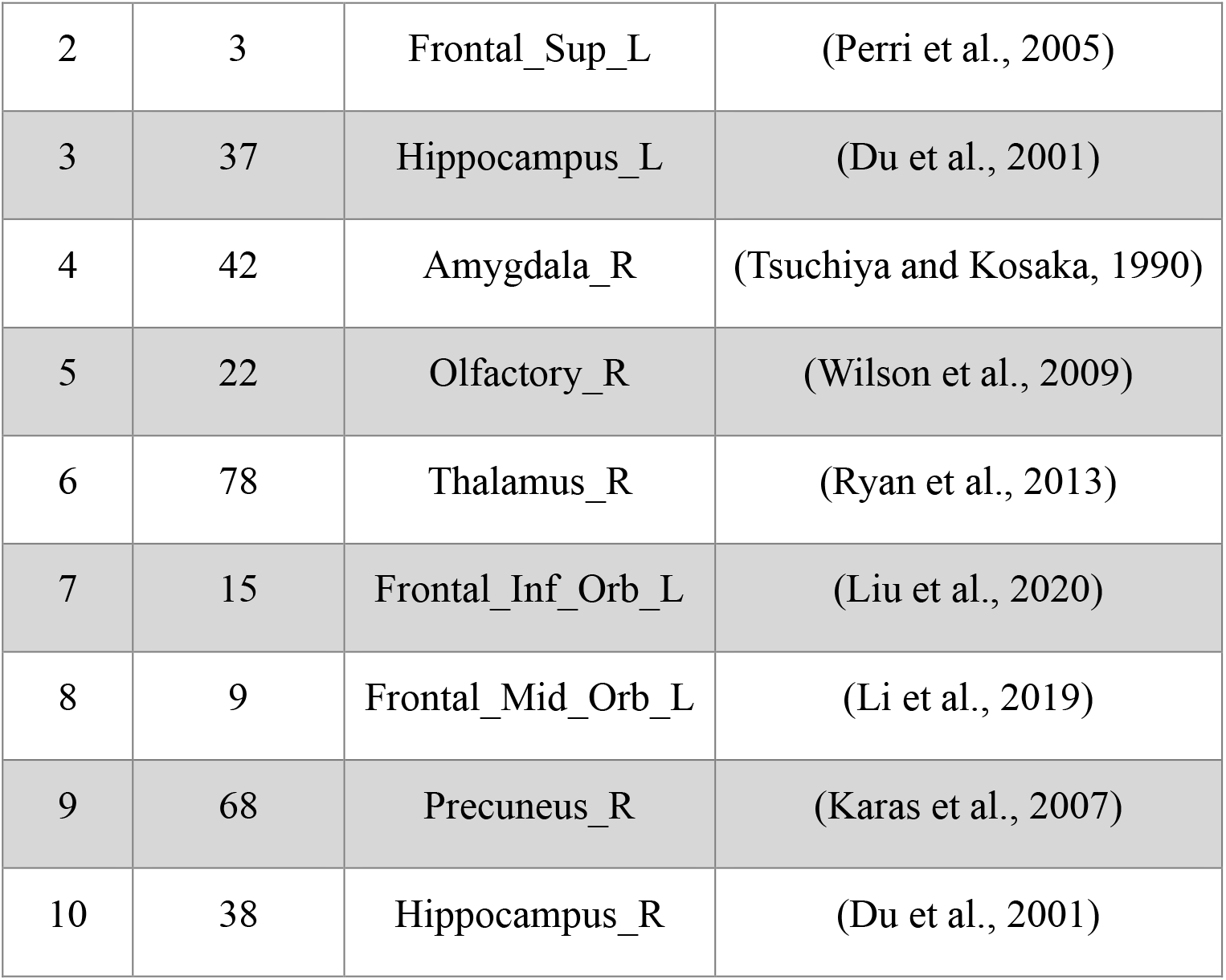
Top ten brain regions in GFS scoring difference between AD and non-AD groups.

Mathematically, we regarded 90 brain regions and its fiber connection as a weighted graph *G*(*V,E, W*), and 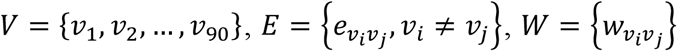, where *v*_*i*_ denotes the *i-*th brain region, 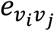 is the edge if there are fibers connection between brain region *v*_*i*_ and *v*_*j*_, and 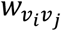 is the edge weight which is the fiber number between the brain region *v*_*i*_ and *v*_*j*_. The average value of the FN matrix of AD is calculated by adding the FN matrix of each AD patient and divided it by the number of AD patients. Similarly, we take the average value of the FN matrix of all normal controls to the FN matrix of the non-AD group.

### 2.2 Local features

#### 2.2.1 Degree Centrality

Let d(*v*_*i*_) denote the degree of a node *v*_*i*_, which is the number of nodes associated with *v*_*i*_. And the degree centrality of a node *v*_*i*_ as follows:

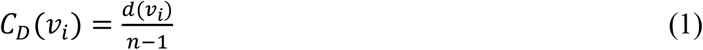

#### 2.2.2 Betweenness centrality

Betweenness centrality c_*B*_ of a node *v*_*i*_ is the sum of the fraction of all-pairs shortest paths that pass through *v*_*i*_:

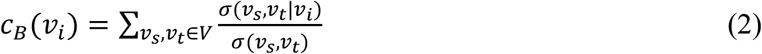

where *σ*(*v*_*s*_, *v*_*t*_) is the number of the shortest paths between *v*_*s*_ and *v*_*t*_, and *σ*(*v*_*s*_, *v*_*s*_|*v*_*i*_) is the number of the shortest paths passing through the node *v*_*i*_. If s *=* t, *σ*(*v*_*s*_, *v*_*s*_) *=* 1, and if *i =* s or *i =* t, *σ*(*v*_*s*_, *v*_*s*_|*v*_*i*_) *=* 0.

In short, if the shortest path between many nodes in the network passes through a point *v*, then *v* has a high degree of betweenness centrality. This node is on the shortcut between other node pairs.

#### 2.2.3 Closeness centrality

Closeness centrality *C*_c_ of a node *v*_*i*_ is the reciprocal of the sum of the shortest path distances from *v*_*i*_ to all n − 1 other nodes. Since the sum of distances depends on the number of nodes in the graph, closeness is normalized by n− 1.

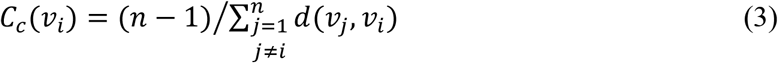

where d(*v*_*j*_, *v*_*i*_) is the shortest path distance between *v*_*j*_ and *v*_*i*_, and nis the number of nodes in the graph.

Closeness centrality is the sum of the distance from a node to all other nodes. The smaller the sum is, the shorter the path from this node to all other nodes is, and the closer the node is to all other nodes. It reflects the proximity between a node and other nodes.

#### 2.2.4 Number of maximal cliques

In graph theory, the clique of graph *G* is a complete subgraph *H* of *G*. *H* is a maximal clique of graph *G* if it is not included by any other clique. The number of maximal cliques of a node can reflect the closeness between the node and other nodes. Only when multiple nodes are all connected can they be considered as maximal cliques. In this paper, we use N_M*C*_(*v*_*i*_) to represent the number of maximal cliques for node *v*_*i*_.

##### 2.2.5 2-hop-connectivity

When examining the correlation of any two nodes in the network, most network analysis methods only consider whether there is an edge connection between two nodes, that is, if there is an edge, the correlation is high, and if there is no connection, the correlation is very weak. In this case, if the edge of the graph is missing due to the disturbance, the result may have a large deviation. For example, for the general random walk (RW) algorithm, the state transition probability is determined by the adjacency matrix of the network. If the adjacency matrix is disturbed, its steady-state probability will change. Generally, when analyzing the correlation of network nodes, the correlation of unconnected nodes in the network will be very low, which is difficult to find the potential characteristics of the network. For any different two nodes in the network, to describe the correlation more accurately, this work not only considers the first-order neighbors between nodes, but also the second-order neighbors between nodes, and we propose an algorithm named 2hopRWR. Finally, each node can get a novel local feature index 2hop-connectivity, which numerical size is represented by local feature score *S*_2−*hop*_. The importance of nodes can be judged based on *S*_2−*hop*_, the larger the *S*_2−*hop*_, the more important the node is.

##### 2.2.5.1 2-hop random walk with restart algorithm

The general random walk on the graph is a transition process by moving from a given node to a randomly selected neighboring node for each step. Consequently, we also regard the node-set {*v*_1_, *v*_**2**_, …, *v*_*n*_} as a set of states {***s***_1_, ***s***_**2**_, …, ***s***_*n*_} in a finite Markov chain ***M***. The transition probability of ***M*** is a conditional probability defined as *P*(*v*_*j*_, *v*_*i*_) *= P****rob***(***s***_***t****+*1_ *= v*_*i*_|***s***_***t***_ *= v*_*j*_) which means that the ***M*** will be at *v*_*i*_ at time ***t*** *+* 1 given that it was at *v*_*j*_ at time ***t. M*** is homogeneous because the transition probability from one state to another is independent of time ***t***. What is more, for any *v*_*j*_ of *V* we have 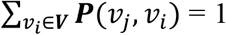. Note that ***M*** is memoryless, therefore, we can define a transition matrix *P ∈ ℝ*^|*V*|×|*V*|^ of ***M***.

Generally, define transition probability *P*(*v*_*j*_, *v*_*i*_) as follows:

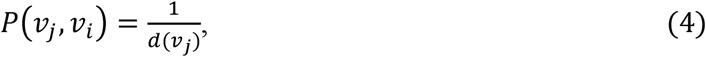

Denote *D*_*G*_ *= diag*{*d*_1_, *d*_**2**_, …, *d*_*n*_} be the diagonal matrix, where 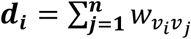 Thus, *P* can be rewritten in matrix notation as follows:

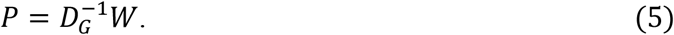

Define *r*_t_ *∈ ℝ*^|*V*|×*1*^ as a vector in which the *i*-th element represents the probability of discovering the random walk at node *v*_*i*_ at step ***t***, so the probability *r*_*t*+1_ can be calculated iteratively by:

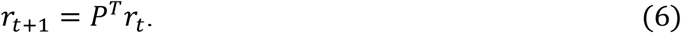

For the random walk with restart (RWR) algorithm (Tong et al., 2006 - 2006), there is an additional restart item compared to the above algorithm. The probability *r*_*t*+1_ can be calculated iteratively by the following expression:

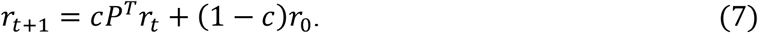

Define initial probability *r*_0_ *∈ ℝ*^|*V*|×1^ as a vector in which the *i*-th element is equal to one, while other elements are zeros. And 1 − *c* is the restart probability (0 ≤ *c* ≤ 1).

But RW and RWR algorithms are all based on the 1-hop neighbor relationship, that is, random walk is based on the existing edge of the graph. If some edges of the graph are missing, the corresponding points cannot be directly transferred, which will lead to a large deviation of the steady-state probability. Therefore, the effectiveness of these algorithms is too much dependent on the integrity of the graph structure. Therefore, in this work, besides the 1-hop neighbor relationship, we also consider the 2-hop neighbors and propose a novel random walk algorithm named 2hopRWR.

The probability *r*_*t*+1_ can be calculated iteratively by:

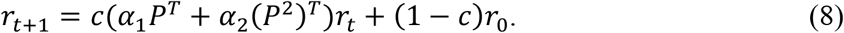

where *α*_1_ and *α*_2_ are the percentage of choosing 1-hop neighbors and 2-hop neighbors, respectively. Specifically, for each point *v*_*i*_ *∈* **V**, *α*_1_ is the ratio of the number of 1-hop neighbors to the total number of 1-hop and 2-hop neighbors, *α*_2_ is the ratio of the number of 2-hop neighbors to the total number of 1-hop and 2-hop neighbors. Therefore, *α*_1_ *+ α*_2_ *=* 1.

At the beginning of the 2hopRWR, we choose a starting node *v*_*i*_, then it would have a probability of c to walk to other nodes and also have a probability of 1 − c to stay in place. Specifically, when the process of walk reaches the node *v*_*j*_, it has a probability of *α*_1_c to walk based on existing edges to 1- hop neighbors and has a probability of *α*_2_c to walk to 2-hop neighbors, and it also has a probability of 1 − c to restart the walk, that is, to go back to the node *v*_*i*_.

After some steps, the RWR will be stable, that is, when ***t*** tends to infinity, *r*_*t*+1_ *= r*_t_. The proof is given in Section 2.2.5.2. When the RWR is stable, stable probability between node *v*_*i*_ and node *v*_*j*_ is defined as the *j*-th element of *r*_t_ corresponding to the starting node is *v*_*i*_.

##### 2.2.5.2 Proof of convergence

Here we will prove that the random walk with restart algorithm is convergent, that is, for equation (8) when ***t*** tends to infinity, *r*_*t*+1_ *= r*_t_.

Define

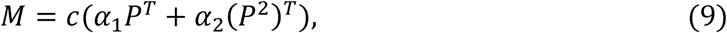

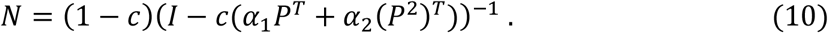

Thus, using (9) and (10) we get

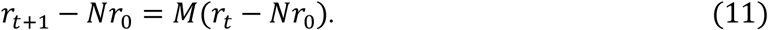

Define

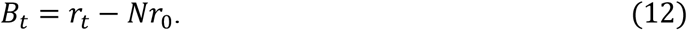

Then

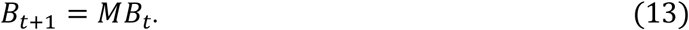

By (13), when t *=* **0**, we have *B*_0_ *=* (**I** − N)*r*_0_, thus

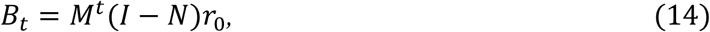

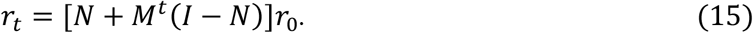

Since 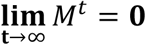 we have

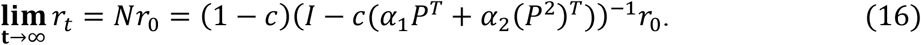

Hence,

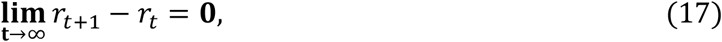

which implies the convergence of the algorithm is proved.

##### 2.2.5.3 hop-connectivity feature score

Any two nodes can be ranked twice according to 2hopRWR. Define П *=* {π_*ij*_}_|*V*|×|*V*|_ be the stable probability matrix where *π*_*ij*_ indicates the stable probability between node *v*_*i*_ and node *v*_*j*_, that is, 2hopRWR starts from node *v*_*i*_ and the probability of reaching node ***v**_j_* when the process is stable. Briefly speaking, the value of *π*_*ij*_ is the *j*-th element of steady-state probability 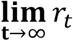 when the *i*-th element of *r*_0_ is 1. Therefore, define a local feature score *S*_2−*hop*_ for node *v*_*i*_ as follows:

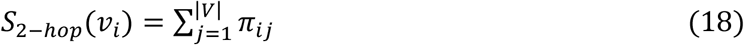

The 2hop-connectivity utilizes the higher-order information of the graph structure relative to the traditional graph theory metrics. Since there is a certain amount of noise in DTI data, a certain threshold value is selected for denoising when DTI data is transformed into brain networks, that is, when the number of fibers between two brain regions is small (less than the threshold value), it is considered that there are no fiber tracts (edges) between two brain regions (nodes). However, since the fiber tract connections between brain regions in Alzheimer’s patients are inherently few, the brain network constructed after denoising may differ more from the real situation. Since 2hop-connectivity is insensitive to 1 hop order relationship and more robust to changes in network structure, 2hop- connectivity is instead more effective for sparse networks (e.g., structural brain networks in Alzheimer’s patients).

### 2.3 Global feature

In this paper, we consider integrating local features to get a new network index: global feature.

We normalized the component of different feature scores to [0, 1]. The normalized features are recorded as *NC*_D_, *NC*_*B*_, *NC*_*C*_, *NN*_M*C*_, and *NS*_2−hop_.

Then, define global feature score *GFS* for node *v*_*i*_ as follows:

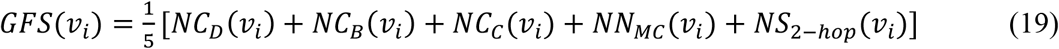

If the *GFS* of a node is relatively high, it means that the node plays a key role in the network.

## 3 Results

### 3.1 Top 10 brain regions for literature verification

By comparing the GFS of the non-AD group with that of the AD group, we got the top ten brain regions in GFS scoring difference (that is, *GFS*_*non*−AD_(*v*_*i*_) − *GFS*_*A*D_(*v*_*i*_)). Then, literature verification was carried out to find out whether these brain regions are related to AD, and the results are shown in Table 1.

For the brain structure network of AD group and non-AD group, the visualization results (Manning et al., 2014) show that the top 10 brain regions are relatively concentrated, as shown in Fig. 4 and Fig. 5.

**FIGURE 1.**
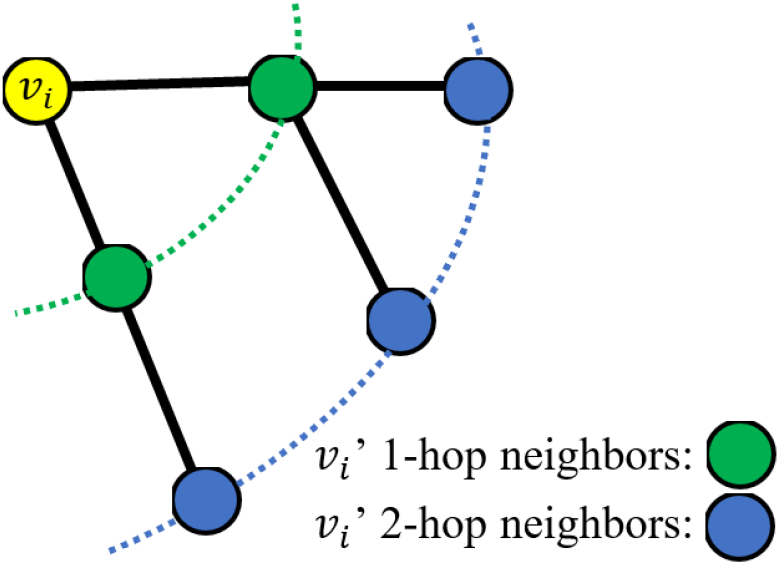
A schematic diagram of 1-hop and 2-hop neighbors

**FIGURE 2.**
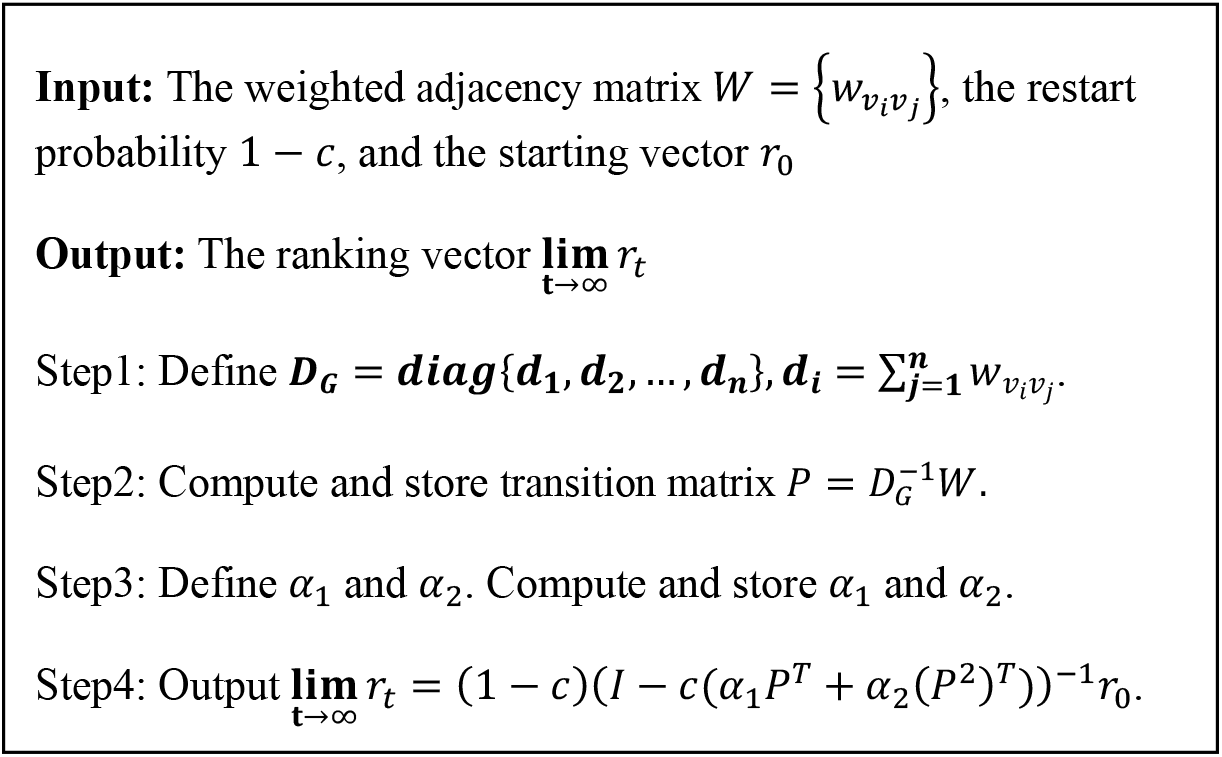
2hopRWR algorithm framework

**FIGURE 3.**
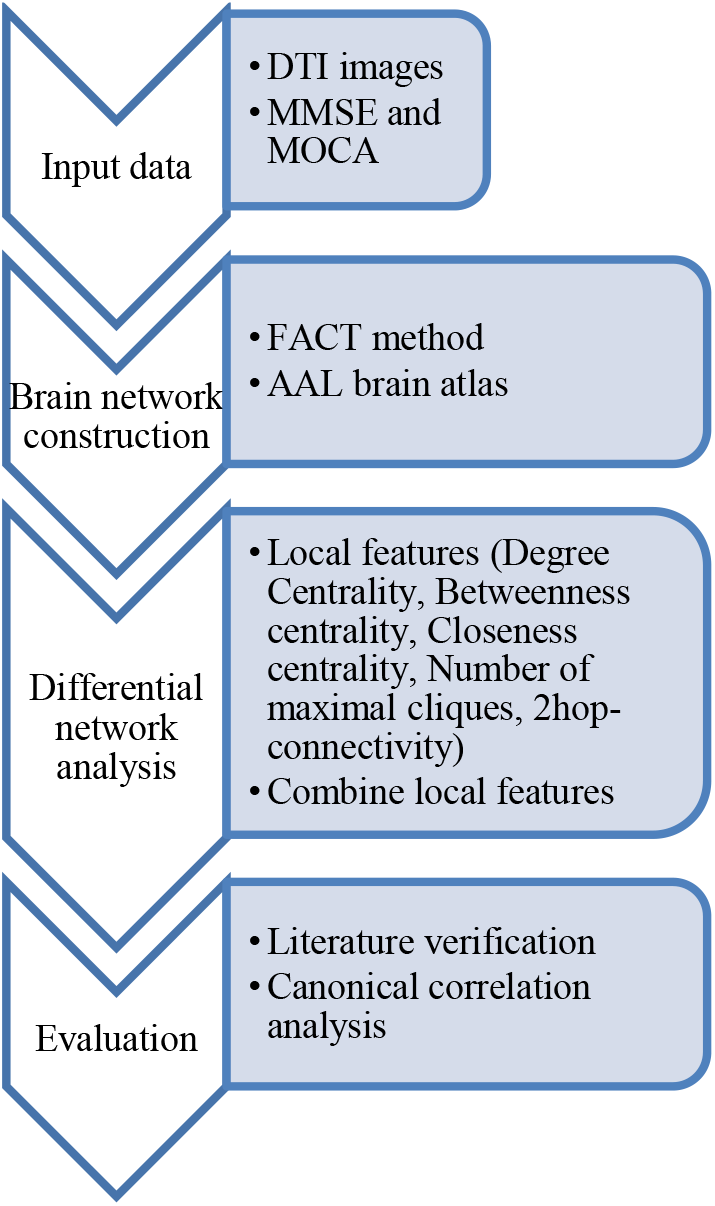
Workflow of our approach.

**FIGURE 4.**
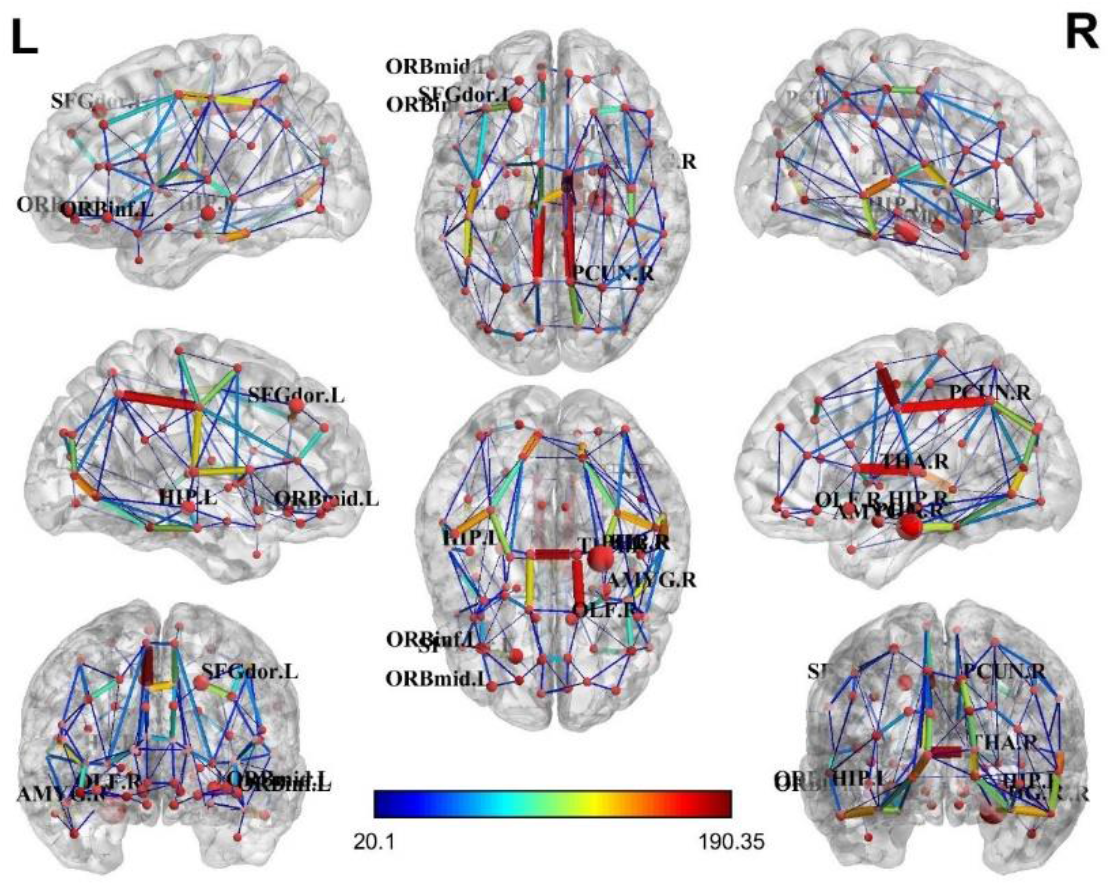
Brain structural network of the AD group.

**FIGURE 5.**
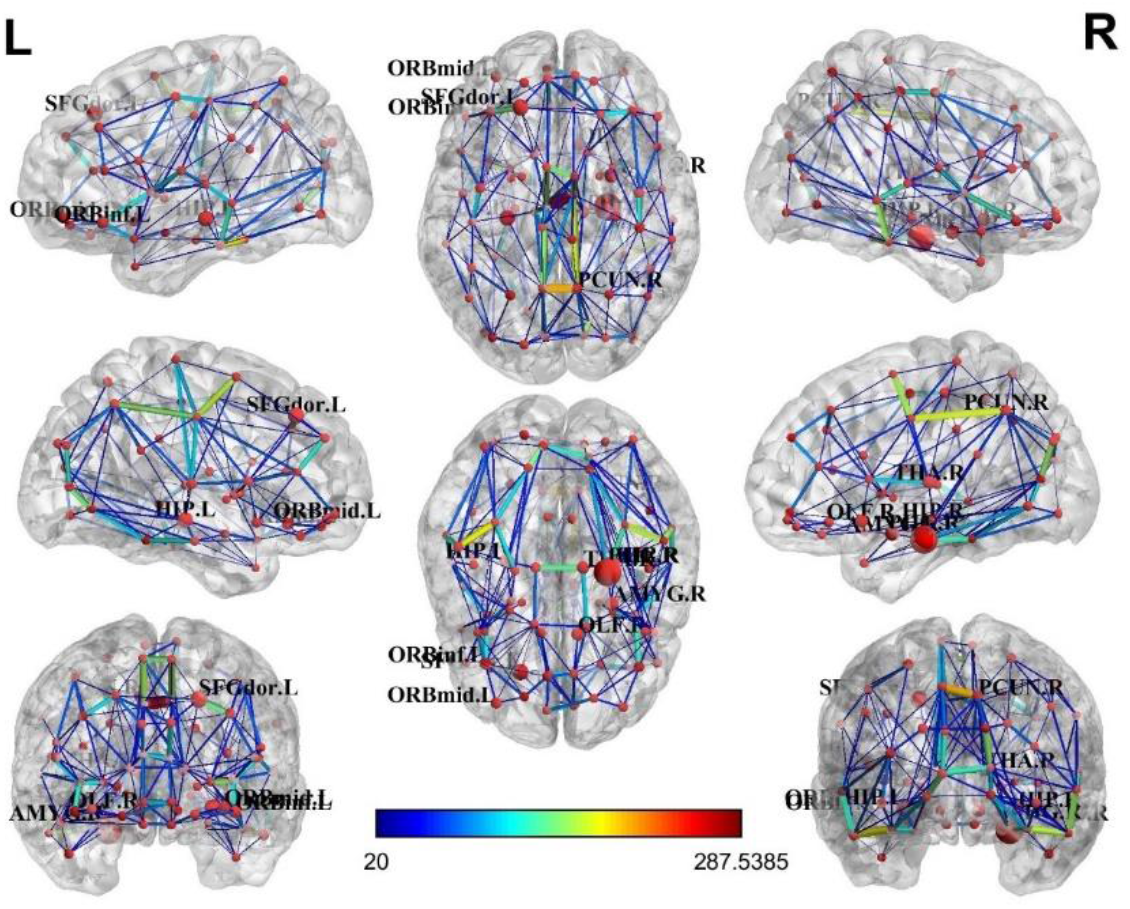
Brain structural network of the non-AD group.

### 3.2 Compare *GFS* with local features

More importantly, we compared the respective predictive effects of all local features. As can be learned from Table 2, whether it is the top 10%, 20%, 30%, or 40% brain regions with the largest differences, the predictive effect of *GFS* is always better than the other local features from the results validated in the literature, as detailed in the **Supplementary Material 4 (SUPPLEMENTARY TABLE 1, SUPPLEMENTARY TABLE 2, and SUPPLEMENTARY TABLE 3)**.

**TABLE 2.** Comparison of the proportion of verified AD-related brain regions for top 10%, 20%, 30%, and 40% ranked by different measures.

|  | NC_D | NC_B | NC_C | NN_MC | NS_2-hop | GFS |
| --- | --- | --- | --- | --- | --- | --- |
| Top 10% | **100.00%** | **100.00%** | **100.00%** | **100.00%** | **100.00%** | **100.00%** |
| Top 20% | 88.89% | 94.44% | 94.44% | 83.33% | 83.33% | **100.00%** |
| Top 30% | 85.19% | 92.59% | 88.89% | 85.19% | 70.37% | **96.30%** |
| Top 40% | 80.56% | 80.56% | **86.11%** | 80.56% | 75.00% | **86.11%** |

### 3.3 Canonical correlation analysis

In this section, we analyzed whether the *GFS* for the Top 10 brain regions is related to the results of MMSE and MoCA. It can analyze whether there is a correlation between two groups of variables. Because some of the people were illiterate, they could not complete the MoCA test, so we took all the people who completed the two scales, a total of 29 people (19 AD and 10 non-AD). At this time, the canonical correlation analysis can be used. Its basic principle is: to grasp the correlation between the two groups of variables as a whole, two composite variables U and V (linear combination of each variable in the two groups) are extracted from the two groups of variables respectively, and the correlation between the two composite variables is used to reflect the overall correlation between the two groups of variables.

The results of the canonical correlation analysis showed that the correlation coefficient between the typical variable pair 1 is 0.7136, which means that there is a very close correlation between *GFS* and MMSE/MOCA scale information.

## 4 Discussion

In this paper, we set a new local feature index 2hop-connectivity to measure the correlation among different areas. The 2hop-connectivity is a node importance metric, like graph-theoretic metrics such as degree centrality. Since most biological networks are relatively sparse, 2hop-connectivity may work better for node importance metrics of sparse networks. And for this, a novel random walk algorithm, 2hopRWR, is proposed, which can compute the local feature index 2hop-connectivity. At the same time, the proof of convergence is given. Next, this paper used the idea of combining 5 local properties to obtain the global feature score (*GFS*), which is more persuasive than using a single network parameter to describe the importance of network nodes. Then, the results of literature verification and canonical correlation analysis also validate the reasonableness and effectiveness of the proposed method. Therefore, *GFS* can be used to distinguish DTI images of the AD group and the non-AD group. Finally, all the top ten brain regions in the *GFS* scoring difference predicted in this paper have been verified by literature. Literature validation results comparing *GFS* with local features showed that *GFS* outperforms individual local features.

Despite the effectiveness of our work, it also has several limitations. First, the mechanism of combining local features is relatively simple. For formula (19), we simply think that the weight of each property is the same, if we can combine more effective information, we can use more reasonable weight distribution to have a deeper understanding of the structure and function network of the brain. Secondly, the brain network constructed in this paper is a structural network, which should be combined with structure and function in the future. For example, if we combine the fMRI data with the existing data to analyze the differences of different brain regions in different tasks, we may get more results.

In brief, *GFS* is expected to be an important and useful index for identifying the difference between network nodes and detecting the changes in information transmission between brain regions in Alzheimer’s disease patients. Moreover, it may provide useful insights into the underlying mechanisms of Alzheimer’s disease. Many older test subjects are illiterate and cannot perform the MMSE and MoCA scales properly, so the *GFS* can be used as a diagnostic aid to infer from DTI imaging data alone whether the subject is likely to be an AD patient, which can greatly reduce the workload of medical workers. Finally, *GFS* can be used as a differential network analysis method (Lichtblau et al., 2017) in other areas of network analysis. We also look forward to the application of the 2hopRWR algorithm in traditional network analysis tasks, such as node classification, link prediction, graph classification.

## 6 Author Contributions

QW designed the work, developed the computing method, wrote the code, analyzed the result, and wrote the manuscript. SC collected clinical data, analyzed the statistics, and polished the manuscript. HW and LC contributed to the imaging work and the statistical analysis. YS designed the work, collected clinical data, and contributed with theoretical support. GY designed the work, analyzed the result, and revised the manuscript.

## 8 Acknowledgments

We thank Prof. Ang Li, Institute of Biophysics, Chinese Academy of Sciences, for his suggestions for the results section.

## 9 Supplementary Material

SUPPLEMENTARY TABLE 1 | MMSE & MoCA data.

SUPPLEMENTARY TABLE 2 | DTI data overview.

SUPPLEMENTARY DATA SHEET 1 | Results of literature validation for *GFS* and local features.

### 1 Data Availability Statement

The code for 2hopRWR in this study can be found in the https://github.com/wangqi27/TwoHopRWR_new.

