## Supplementary Material 4 for "Predicting Brain Regions Related to Alzheimer’s Disease Based on Global Feature"

**SUPPLEMENTARY TABLE 1** | Literature validation results of brain regions related to Alzheimer's Disease (AD) in AAL template (90 regions). In particular, the relevant kinds of the literature of region ID 77, 78, 81, 82, 85, and 86 were found based on literature (Zhang et al., 2015)

| Region ID | AAL regions | Abbreviation | Evidence |
|---|---|---|---|
| 1 | Precental gyrus | PreCG.L | (Kang et al., 2013b) |
| 2 | Precental gyrus | PreCG.R | No literature was found |
| 3 | Superior frontal gyrus, dorsolateral | SFGdor.L | (Qin et al., 2019) |
| 4 | Superior frontal gyrus, dorsolateral | SFGdor.R | No literature was found |
| 5 | Superior frontal gyrus, orbital part | ORBsup.L | No literature was found |
| 6 | Superior frontal gyrus, orbital part | ORBsup.R | No literature was found |
| 7 | Middle frontal gyrus | MFG.L | (Schultz et al., 2015) |
| 8 | Middle frontal gyrus | MFG.R | (Schultz et al., 2015) |
| 9 | Middle frontal gyrus, orbital part | ORBmid.L | (Li et al., 2019) |
| 10 | Middle frontal gyrus, orbital part | ORBmid.R | (Li et al., 2019) |
| 11 | Inferior frontal gyrus, opercular part | IFGoperc.L | No literature was found |
| 12 | Inferior frontal gyrus, opercular part | IFGoperc.R | No literature was found |
| 13 | Inferior frontal gyrus, triangular part | IFGtriang.L | No literature was found |
| 14 | Inferior frontal gyrus, triangular part | IFGtriang.R | No literature was found |
| 15 | Inferior frontal gyrus, orbital part | ORBinf.L | (Liu et al., 2020) |
| 16 | Inferior frontal gyrus, orbital part | ORBinf.R | (Shan et al., 2018) |
| 17 | Rolandic operculum | ROL.L | No literature was found |
| 18 | Rolandic operculum | ROL.R | No literature was found |
| 19 | Supplementary motor area | SMA.L | No literature was found |
| 20 | Supplementary motor area | SMA.R | No literature was found |
| 21 | Olfactory cortex | OLF.L | (Reyes et al., 1986) |
| 22 | Olfactory cortex | OLF.R | (Reyes et al., 1986) |
| 23 | Superior frontal gyrus, medial | SFGmed.L | (Hallam et al., 2020) |
| 24 | Superior frontal gyrus, medial | SFGmed.R | (Hallam et al., 2020) |
| 25 | Superior frontal gyrus, medial orbital | ORBsupmed.L | (Hallam et al., 2020) |
| 26 | Superior frontal gyrus, medial orbital | ORBsupmed.R | (Hallam et al., 2020) |
| 27 | Gyrus rectus | REC.L | (Mölsä et al., 1987) |
| 28 | Gyrus rectus | REC.R | (Mölsä et al., 1987) |
| 29 | Insula | INS.L | (Foundas et al., 1997) |
| 30 | Insula | INS.R | (Foundas et al., 1997) |
| 31 | Anterior cingulate and paracingulate gyri | ACG.L | No literature was found |
| 32 | Anterior cingulate and paracingulate gyri | ACG.R | No literature was found |
| 33 | Median cingulate and paracingulate gyri | DCG.L | No literature was found |
| 34 | Median cingulate and paracingulate gyri | DCG.R | No literature was found |
| 35 | Posterior cingulate gyrus | PCG.L | (Scheff et al., 2015) |
| 36 | Posterior cingulate gyrus | PCG.R | (Scheff et al., 2015) |
| 37 | Hippocampus | HIP.L | (Foundas et al., 1997) |
| 38 | Hippocampus | HIP.R | (Foundas et al., 1997) |

| | | | |
|---|---|---|---|
| 39 | Parahippocampal gyrus | PHG.L | (Eskildsen et al., 2015) |
| 40 | Parahippocampal gyrus | PHG.R | (Eskildsen et al., 2015) |
| 41 | Amygdala | AMYG.L | (Tsuchiya and Kosaka, 1990) |
| 42 | Amygdala | AMYG.R | (Tsuchiya and Kosaka, 1990) |
| 43 | Calcarine fissure and surrounding cortex | CAL.L | (Ren et al., 2020) |
| 44 | Calcarine fissure and surrounding cortex | CAL.R | (Ren et al., 2020) |
| 45 | Cuneus | CUN.L | No literature was found |
| 46 | Cuneus | CUN.R | No literature was found |
| 47 | Lingual gyrus | LING.L | (Lehmann et al., 2013) |
| 48 | Lingual gyrus | LING.R | (Lehmann et al., 2013) |
| 49 | Superior occipital gyrus | SOG.L | (Beyer et al., 2012) |
| 50 | Superior occipital gyrus | SOG.R | (Beyer et al., 2012) |
| 51 | Middle occipital gyrus | MOG.L | (Lehmann et al., 2013) |
| 52 | Middle occipital gyrus | MOG.R | (Lehmann et al., 2013) |
| 53 | Inferior occipital gyrus | IOG.L | (Liu et al., 2018) |
| 54 | Inferior occipital gyrus | IOG.R | No literature was found |
| 55 | Fusiform gyrus | FFG.L | (Ma et al., 2020) |
| 56 | Fusiform gyrus | FFG.R | (Ma et al., 2020) |
| 57 | Postcentral gyrus | PoCG.L | (Kang et al., 2013b) |
| 58 | Postcentral gyrus | PoCG.R | (Kang et al., 2013b) |
| 59 | Superior parietal gyrus | SPG.L | (Vasconcelos et al., 2014) |
| 60 | Superior parietal gyrus | SPG.R | (Vasconcelos et al., 2014) |
| 61 | Inferior parietal, but supramarginal and angular gyri | IPL.L | No literature was found |
| 62 | Inferior parietal, but supramarginal and angular gyri | IPL.R | No literature was found |
| 63 | Supramarginal gyrus | SMG.L | (Grignon et al., 1998) |
| 64 | Supramarginal gyrus | SMG.R | (Grignon et al., 1998) |
| 65 | Angular gyrus | ANG.L | No literature was found |
| 66 | Angular gyrus | ANG.R | No literature was found |
| 67 | Precuneus | PCUN.L | (Karas et al., 2007) |
| 68 | Precuneus | PCUN.R | (Karas et al., 2007) |
| 69 | Paracentral lobule | PCL.L | (Kang et al., 2013a) |
| 70 | Paracentral lobule | PCL.R | (Kang et al., 2013a) |
| 71 | Caudate nucleus | CAU.L | (Möller et al., 2015) |
| 72 | Caudate nucleus | CAU.R | (Möller et al., 2015) |
| 73 | Lenticular nucleus, putamen | PUT.L | No literature was found |
| 74 | Lenticular nucleus, putamen | PUT.R | No literature was found |
| 75 | Lenticular nucleus, pallidum | PAL.L | No literature was found |

| | | | |
|---|---|---|---|
| 76 | Lenticular nucleus, pallidum | PAL.R | No literature was found |
| 77 | Thalamus | THA.L | (He et al., 2015) |
| 78 | Thalamus | THA.R | (He et al., 2015) |
| 79 | Heschl gyrus | HES.L | No literature was found |
| 80 | Heschl gyrus | HES.R | No literature was found |
| 81 | Superior temporal gyrus | STG.L | (Paakki et al., 2010) |
| 82 | Superior temporal gyrus | STG.R | (Paakki et al., 2010) |
| 83 | Temporal pole: superior temporal gyrus | TPOsup.L | No literature was found |
| 84 | Temporal pole: superior temporal gyrus | TPOsup.R | No literature was found |
| 85 | Middle temporal gyrus | MTG.L | (Aubry et al., 2015) |
| 86 | Middle temporal gyrus | MTG.R | (Aubry et al., 2015) |
| 87 | Temporal pole: middle temporal gyrus | TPOmid.L | (Sturm et al., 2013) |
| 88 | Temporal pole: middle temporal gyrus | TPOmid.R | No literature was found |
| 89 | Inferior temporal gyrus | ITG.L | (Scheff et al., 2011) |
| 90 | Inferior temporal gyrus | ITG.R | (Scheff et al., 2011) |

**SUPPLEMENTARY TABLE 2|** Ranking results of different measures for AAL template (90 regions). The filling color of yellow indicates that the brain area has been verified to be related to AD.

| NC_B | NC_C | NN_MC | NS_2hop | NC_D | GFS |
|---|---|---|---|---|---|
| 78 | 3 | 37 | 42 | 3 | 40 |
| 37 | 40 | 40 | 3 | 9 | 3 |
| 40 | 9 | 38 | 40 | 22 | 37 |
| 29 | 37 | 78 | 22 | 40 | 42 |
| 68 | 78 | 77 | 49 | 15 | 22 |
| 42 | 22 | 15 | 16 | 42 | 78 |
| 3 | 42 | 42 | 58 | 37 | 15 |
| 58 | 15 | 68 | 15 | 59 | 9 |
| 1 | 89 | 67 | 7 | 78 | 68 |
| 77 | 58 | 73 | 45 | 89 | 38 |
| 7 | 59 | 71 | 23 | 16 | 77 |
| 59 | 68 | 22 | 2 | 58 | 58 |
| 38 | 16 | 48 | 8 | 21 | 16 |
| 89 | 38 | 31 | 10 | 1 | 29 |
| 84 | 29 | 29 | 50 | 14 | 89 |
| 60 | 77 | 3 | 1 | 29 | 59 |
| 8 | 1 | 72 | 9 | 53 | 1 |
| 22 | 84 | 5 | 13 | 79 | 7 |
| 16 | 7 | 21 | 59 | 7 | 49 |
| 43 | 21 | 6 | 17 | 8 | 84 |
| 15 | 60 | 89 | 14 | 10 | 21 |
| 9 | 14 | 82 | 62 | 27 | 8 |
| 82 | 53 | 27 | 5 | 60 | 27 |
| 79 | 27 | 9 | 27 | 38 | 23 |
| 53 | 49 | 58 | 6 | 68 | 43 |
| 30 | 79 | 30 | 60 | 84 | 67 |
| 39 | 8 | 43 | 51 | 5 | 71 |
| 51 | 10 | 84 | 43 | 77 | 2 |
| 2 | 23 | 16 | 53 | 49 | 5 |
| 80 | 71 | 49 | 67 | 2 | 60 |
| 88 | 43 | 47 | 70 | 23 | 10 |
| 23 | 2 | 28 | 37 | 80 | 14 |
| 14 | 5 | 32 | 71 | 43 | 48 |
| 54 | 30 | 1 | 68 | 51 | 53 |
| 49 | 48 | 88 | 28 | 17 | 79 |
| 10 | 51 | 23 | 4 | 30 | 30 |
| 27 | 67 | 25 | 69 | 88 | 51 |
| 24 | 80 | 33 | 21 | 13 | 88 |
| 21 | 88 | 2 | 63 | 48 | 6 |

| | | | | | |
|---|---|---|---|---|---|
| 12 | 13 | 81 | 19 | 71 | 80 |
| 70 | 39 | 19 | 11 | 39 | 17 |
| 55 | 17 | 4 | 79 | 25 | 13 |
| 48 | 25 | 10 | 41 | 24 | 39 |
| 17 | 82 | 39 | 89 | 67 | 28 |
| 45 | 47 | 14 | 77 | 6 | 47 |
| 65 | 24 | 24 | 85 | 28 | 24 |
| 44 | 6 | 51 | 81 | 47 | 25 |
| 67 | 28 | 46 | 48 | 82 | 45 |
| 5 | 55 | 59 | 57 | 55 | 73 |
| 36 | 46 | 7 | 61 | 4 | 55 |
| 19 | 4 | 86 | 75 | 46 | 82 |
| 47 | 61 | 55 | 47 | 54 | 19 |
| 18 | 54 | 41 | 87 | 26 | 70 |
| 66 | 26 | 60 | 65 | 44 | 46 |
| 62 | 44 | 8 | 24 | 45 | 61 |
| 71 | 45 | 17 | 55 | 61 | 44 |
| 25 | 70 | 79 | 83 | 73 | 4 |
| 28 | 73 | 26 | 84 | 70 | 62 |
| 35 | 65 | 13 | 88 | 12 | 54 |
| 46 | 12 | 53 | 44 | 19 | 50 |
| 13 | 19 | 75 | 25 | 41 | 65 |
| 61 | 62 | 80 | 30 | 86 | 12 |
| 69 | 41 | 61 | 80 | 81 | 26 |
| 6 | 86 | 45 | 73 | 62 | 81 |
| 86 | 81 | 54 | 12 | 72 | 41 |
| 64 | 32 | 50 | 29 | 32 | 86 |
| 26 | 50 | 90 | 35 | 50 | 32 |
| 63 | 75 | 69 | 32 | 65 | 75 |
| 50 | 57 | 44 | 46 | 83 | 69 |
| 11 | 72 | 57 | 38 | 33 | 57 |
| 52 | 83 | 65 | 39 | 57 | 66 |
| 75 | 36 | 70 | 64 | 75 | 35 |
| 41 | 35 | 63 | 26 | 66 | 72 |
| 73 | 85 | 62 | 90 | 36 | 85 |
| 81 | 66 | 36 | 66 | 35 | 83 |
| 32 | 33 | 52 | 86 | 85 | 36 |
| 85 | 64 | 66 | 31 | 31 | 64 |
| 87 | 69 | 64 | 33 | 18 | 11 |
| 57 | 11 | 85 | 54 | 11 | 63 |
| 20 | 18 | 12 | 78 | 64 | 31 |
| 76 | 31 | 76 | 20 | 69 | 33 |
| 4 | 52 | 87 | 72 | 52 | 18 |

| | | | | | |
|---|---|---|---|---|---|
| 83 | 63 | 18 | 52 | 63 | 52 |
| 90 | 90 | 83 | 36 | 90 | 90 |
| 34 | 20 | 35 | 76 | 76 | 87 |
| 33 | 76 | 56 | 56 | 56 | 76 |
| 72 | 56 | 20 | 18 | 20 | 20 |
| 56 | 87 | 11 | 82 | 34 | 56 |
| 31 | 34 | 74 | 34 | 87 | 34 |
| 74 | 74 | 34 | 74 | 74 | 74 |

**SUPPLEMENTARY TABLE 3 |** Comparison of the proportion of verified AD-related brain regions by different measures.

| Rank | NC_B | NC_C | NN_MC | NS_2hop | NC_D | GFS |
|---|---|---|---|---|---|---|
| 1 | 100.00% | 100.00% | 100.00% | 100.00% | 100.00% | 100.00% |
| 2 | 100.00% | 100.00% | 100.00% | 100.00% | 100.00% | 100.00% |
| 3 | 100.00% | 100.00% | 100.00% | 100.00% | 100.00% | 100.00% |
| 4 | 100.00% | 100.00% | 100.00% | 100.00% | 100.00% | 100.00% |
| 5 | 100.00% | 100.00% | 100.00% | 100.00% | 100.00% | 100.00% |
| 6 | 100.00% | 100.00% | 100.00% | 100.00% | 100.00% | 100.00% |
| 7 | 100.00% | 100.00% | 100.00% | 100.00% | 100.00% | 100.00% |
| 8 | 100.00% | 100.00% | 100.00% | 100.00% | 100.00% | 100.00% |
| 9 | 100.00% | 100.00% | 100.00% | 100.00% | 100.00% | 100.00% |
| 10 | 100.00% | 100.00% | 90.00% | 90.00% | 100.00% | 100.00% |
| 11 | 100.00% | 100.00% | 90.91% | 90.91% | 100.00% | 100.00% |
| 12 | 100.00% | 100.00% | 91.67% | 83.33% | 100.00% | 100.00% |
| 13 | 100.00% | 100.00% | 92.31% | 84.62% | 100.00% | 100.00% |
| 14 | 100.00% | 100.00% | 85.71% | 85.71% | 100.00% | 100.00% |
| 15 | 93.33% | 100.00% | 86.67% | 86.67% | 93.33% | 100.00% |
| 16 | 93.75% | 100.00% | 87.50% | 87.50% | 93.75% | 100.00% |
| 17 | 94.12% | 100.00% | 88.24% | 88.24% | 94.12% | 100.00% |
| 18 | 94.44% | 94.44% | 83.33% | 83.33% | 88.89% | 100.00% |
| 19 | 94.74% | 94.74% | 84.21% | 84.21% | 89.47% | 100.00% |
| 20 | 95.00% | 95.00% | 80.00% | 80.00% | 90.00% | 95.00% |
| 21 | 95.24% | 95.24% | 80.95% | 76.19% | 90.48% | 95.24% |
| 22 | 95.45% | 90.91% | 81.82% | 72.73% | 90.91% | 95.45% |
| 23 | 95.65% | 91.30% | 82.61% | 69.57% | 91.30% | 95.65% |
| 24 | 91.67% | 91.67% | 83.33% | 70.83% | 91.67% | 95.83% |
| 25 | 92.00% | 92.00% | 84.00% | 68.00% | 92.00% | 96.00% |
| 26 | 92.31% | 88.46% | 84.62% | 69.23% | 88.46% | 96.15% |
| 27 | 92.59% | 88.89% | 85.19% | 70.37% | 85.19% | 96.30% |
| 28 | 92.86% | 89.29% | 82.14% | 71.43% | 85.71% | 92.86% |
| 29 | 89.66% | 89.66% | 82.76% | 72.41% | 86.21% | 89.66% |
| 30 | 86.67% | 90.00% | 83.33% | 73.33% | 83.33% | 90.00% |
| 31 | 83.87% | 90.32% | 83.87% | 74.19% | 83.87% | 90.32% |
| 32 | 84.38% | 87.50% | 84.38% | 75.00% | 81.25% | 87.50% |
| 33 | 81.82% | 84.85% | 81.82% | 75.76% | 81.82% | 87.88% |
| 34 | 79.41% | 85.29% | 82.35% | 76.47% | 82.35% | 88.24% |
| 35 | 80.00% | 85.71% | 80.00% | 77.14% | 80.00% | 85.71% |
| 36 | 80.56% | 86.11% | 80.56% | 75.00% | 80.56% | 86.11% |
| 37 | 81.08% | 86.49% | 81.08% | 75.68% | 78.38% | 86.49% |
| 38 | 81.58% | 84.21% | 78.95% | 76.32% | 76.32% | 84.21% |
| 39 | 82.05% | 82.05% | 76.92% | 76.92% | 76.92% | 82.05% |
| 40 | 80.00% | 80.00% | 77.50% | 75.00% | 77.50% | 80.00% |

| | | | | | | |
|---|---|---|---|---|---|---|
| 41 | 80.49% | 80.49% | 75.61% | 73.17% | 78.05% | 78.05% |
| 42 | 80.95% | 78.57% | 73.81% | 71.43% | 78.57% | 76.19% |
| 43 | 81.40% | 79.07% | 74.42% | 72.09% | 79.07% | 76.74% |
| 44 | 79.55% | 79.55% | 75.00% | 72.73% | 79.55% | 77.27% |
| 45 | 77.78% | 80.00% | 73.33% | 73.33% | 77.78% | 77.78% |
| 46 | 76.09% | 80.43% | 73.91% | 73.91% | 78.26% | 78.26% |
| 47 | 76.60% | 78.72% | 74.47% | 74.47% | 78.72% | 78.72% |
| 48 | 77.08% | 79.17% | 72.92% | 75.00% | 79.17% | 77.08% |
| 49 | 75.51% | 79.59% | 73.47% | 75.51% | 79.59% | 75.51% |
| 50 | 76.00% | 78.00% | 74.00% | 74.00% | 78.00% | 76.00% |
| 51 | 74.51% | 76.47% | 74.51% | 72.55% | 76.47% | 76.47% |
| 52 | 75.00% | 75.00% | 75.00% | 73.08% | 75.00% | 75.00% |
| 53 | 73.58% | 73.58% | 75.47% | 73.58% | 75.47% | 75.47% |
| 54 | 72.22% | 74.07% | 75.93% | 72.22% | 75.93% | 74.07% |
| 55 | 70.91% | 74.55% | 76.36% | 72.73% | 74.55% | 72.73% |
| 56 | 71.43% | 73.21% | 75.00% | 73.21% | 73.21% | 73.21% |
| 57 | 71.93% | 73.68% | 73.68% | 71.93% | 71.93% | 71.93% |
| 58 | 72.41% | 72.41% | 74.14% | 70.69% | 72.41% | 70.69% |
| 59 | 72.88% | 71.19% | 72.88% | 69.49% | 71.19% | 69.49% |
| 60 | 71.67% | 70.00% | 73.33% | 70.00% | 70.00% | 70.00% |
| 61 | 70.49% | 68.85% | 72.13% | 70.49% | 70.49% | 68.85% |
| 62 | 69.35% | 67.74% | 70.97% | 70.97% | 70.97% | 67.74% |
| 63 | 69.84% | 68.25% | 69.84% | 69.84% | 71.43% | 68.25% |
| 64 | 68.75% | 68.75% | 68.75% | 68.75% | 70.31% | 68.75% |
| 65 | 69.23% | 69.23% | 67.69% | 67.69% | 70.77% | 69.23% |
| 66 | 69.70% | 68.18% | 68.18% | 68.18% | 69.70% | 69.70% |
| 67 | 70.15% | 68.66% | 68.66% | 68.66% | 70.15% | 68.66% |
| 68 | 70.59% | 67.65% | 69.12% | 67.65% | 69.12% | 67.65% |
| 69 | 71.01% | 68.12% | 69.57% | 66.67% | 68.12% | 68.12% |
| 70 | 70.00% | 68.57% | 70.00% | 67.14% | 67.14% | 68.57% |
| 71 | 70.42% | 67.61% | 69.01% | 67.61% | 67.61% | 67.61% |
| 72 | 69.44% | 68.06% | 69.44% | 68.06% | 66.67% | 68.06% |
| 73 | 69.86% | 68.49% | 69.86% | 68.49% | 65.75% | 68.49% |
| 74 | 68.92% | 68.92% | 68.92% | 68.92% | 66.22% | 68.92% |
| 75 | 69.33% | 68.00% | 69.33% | 68.00% | 66.67% | 68.00% |
| 76 | 68.42% | 67.11% | 69.74% | 68.42% | 67.11% | 68.42% |
| 77 | 68.83% | 67.53% | 68.83% | 67.53% | 66.23% | 68.83% |
| 78 | 69.23% | 67.95% | 69.23% | 66.67% | 65.38% | 67.95% |
| 79 | 69.62% | 67.09% | 69.62% | 65.82% | 64.56% | 68.35% |
| 80 | 68.75% | 66.25% | 68.75% | 66.25% | 65.00% | 67.50% |
| 81 | 67.90% | 65.43% | 67.90% | 65.43% | 65.43% | 66.67% |
| 82 | 67.07% | 65.85% | 68.29% | 65.85% | 65.85% | 65.85% |
| 83 | 66.27% | 66.27% | 67.47% | 66.27% | 66.27% | 66.27% |

| | | | | | | |
|---|---|---|---|---|---|---|
| 84 | 66.67% | 66.67% | 66.67% | 66.67% | 66.67% | 66.67% |
| 85 | 65.88% | 65.88% | 67.06% | 65.88% | 65.88% | 67.06% |
| 86 | 65.12% | 65.12% | 67.44% | 66.28% | 66.28% | 66.28% |
| 87 | 65.52% | 65.52% | 66.67% | 65.52% | 65.52% | 65.52% |
| 88 | 65.91% | 65.91% | 65.91% | 65.91% | 64.77% | 65.91% |
| 89 | 65.17% | 65.17% | 65.17% | 65.17% | 65.17% | 65.17% |
| 90 | 64.44% | 64.44% | 64.44% | 64.44% | 64.44% | 64.44% |
